# Long shelf-life streptavidin support-films suitable for electron microscopy of biological macromolecules

**DOI:** 10.1101/054452

**Authors:** Bong-Gyoon Han, Zoe Watson, Hannah Kang, Arto Pulk, Kenneth H. Downing, Jamie Cate, Robert M. Glaeser

## Abstract

We describe a rapid and convenient method of growing streptavidin (SA) monolayer crystals directly on holey-carbon EM grids. As expected, these SA monolayer crystals retain their biotin-binding function and crystalline order through a cycle of embedding in trehalose and, later, its removal. This fact allows one to prepare, and store for later use, EM grids on which SA monolayer crystals serve as an affinity substrate for preparing specimens of biological macromolecules. In addition, we report that coating the lipid-tail side of trehalose-embedded monolayer crystals with evaporated carbon appears to improve the consistency with which well-ordered, single crystals are observed to span over entire, 2 μm holes of the support films. Randomly biotinylated 70 S ribosomes are used as a test specimen to show that these support films can be used to obtain a high-resolution cryo-EM structure.

## INTRODUCTION

There are a number of reasons to consider using streptavidin monolayer crystals as “affinity” support films for cryo-electron microscopy (cryo-EM). Macromolecules of interest can be easily tagged with biotin or a streptavidin-binding peptide and then bound to streptavidin (SA) with high affinity and specificity. Furthermore, tagging followed by affinity binding is expected to pose less risk to the native structure of the macromolecule than does (1) adsorption of particles to the surface of carbon film, even when rendered hydrophilic by exposure to a glow discharge, or (2) repeated collision with the air-water interface that occurs when freely diffusing macromolecules are confined to a thin aqueous film (Taylor and Glaeser, 2008).

Monolayer crystals of SA have been considered previously by several authors for use as an affinity-support film. One early study viewed SA as being a “general adaptor” for linking any kind of biotinylated molecule to a lipid monolayer (Darst et al., 1991). Chemically biotinylated ferritin was used in that work to show that a high density of randomly distributed particles could be bound to 2-D crystals of SA. In an extension of the adaptor-molecule idea, Crucifix et al. first randomly decorated SA monolayer crystals with biotinylated dsDNA molecules, and then used the immobilized DNA as bait to bind yeast RNA Pol I particles (Crucifix et al., 2004). Wang et al. showed that biotinylated proteoliposomes could be bound at high density (Wang and Sigworth, 2009; Wang et al., 2008), and they introduced the further innovation of eliminating the periodic background due to SA by masking out the Bragg reflections in the computed Fourier transforms of images. Han et al. then went on to demonstrate the generality with which chemical biotinylation of soluble-protein complexes could be used (Han et al., 2012).

In spite of these promising demonstrations, SA monolayer crystals have not been adopted as support films for routine data collection. Our recent attempts to do so made it clear that two major problems remained. (1) The standard protocol for growing monolayer crystals involves an additional incubation step of 2 hours or more (Wang and Sigworth, 2010), which both slows and complicates the process of preparing cryo-EM specimens. (2) In addition, while the results can be excellent, the growth of large, well-ordered crystals on micro-wells, together with their transfer onto EM grids, is quite inconsistent.

We now describe a simplified, on-grid crystallization protocol that yields large SA crystals in times as short as 10 minutes. In addition, we demonstrate that trehalose-embedding makes it possible to prepare these grids in advance, with their useful shelf life expected to be months or longer. We also find that a thin layer of evaporated carbon can be deposited on the back side (lipid-tail side) of the trehalose-embedded SA crystals in order to add mechanical stability. In a practical test of these grids, *E. coli* 70S ribosomes were used to obtain a 3-D reconstruction at a global resolution estimated to be ~4.0 Å, which improved to ~3.9 Å when focused refinement was used for the large subunit.

## MATERIALS AND METHODS

### Lipids

The biotinylated lipid used here is 1,2-dipalmitoyl-sn-glycero-3-phosphoethanolamine-N-(cap biotinyl), supplied as a 10 mg/mL solution in chloroform/methanol/water (Avanti Polar Lipids). This was diluted to 1.0 mg/mL with a solution of chloroform/methanol/water and aliquoted into small volumes intended for a single usage. The aliquots were sealed under nirogen gas and stored at −80 °C. No deterioration as a function of time was observed in the ability of such aliquots to produce high-quality streptavidin monolayer crystals. Nevertheless, as a precaution, we prepare new aliquots after a period of about 6 months.

### Streptavidin

Streptavidin (SA) was purchased from New England Biolabs (catalog number N7021S). This sample is provided at a concentration of ~1 mg/mL, dissolved in 10 mM sodium phosphate pH 7.2 with 0.15 M NaCl. This was aliquoted in quantities intended for single use, frozen in liquid nitrogen, and stored at −80 °C. Similar to what we do for the lipid, as a precaution, we prepare new aliquots of streptavidin after a period of about 6 months.

### Protocol for growing monolayer crystals directly on holey-carbon EM grids

A lipid monolayer, cast on an air-water interface, is first picked up by touching the lipid from above with a hydrophobic, holey-carbon EM grid. This results in Langmuir-Schaefer transfer of patches of the monolayer that span the holes of the carbon film, as was discovered by (Kubalek et al., 1991). We presume that an additional, unwanted lipid monolayer is also transferred to the air-water surface of the small volume of buffer that adheres to the (now hydrophilic) face of the EM grid. Thus, to remove as much lipid as possible from the surface of the adhering drop, we touch the grid to three successive, 50 μL drops of subphase buffer sitting on parafilm. Next, SA is added to the small droplet of buffer that adheres to the face of the EM grid. The grid is then incubated long enough to allow binding and subsequent crystallization of SA. We refer to this technique as the “on-grid” crystallization method. Figure S1 shows two photographs that further illustrate the steps just described.

We cast the above-mentioned lipid monolayer on a small trough holding ~5 mL of subphase buffer consisting of 50 mM HEPES (pH 7.5), 0.15 M KCl, and 10 % trehalose (Swanson Health Products). Including trehalose in the subphase buffer serves only to eliminate the need for a buffer-exchange “wash” before air-drying (see the following section). This simplification became possible after we observed that adding 10% trehalose has no effect on the quality of monolayer crystals of streptavidin. Before applying lipid, however, we first “dust” the air-water interface lightly with unscented talcum powder, and then we apply a small droplet (~10 µL) of castor oil at the center. As the castor oil spreads, the advancing front of oil sweeps away contaminants at the air-water interface and compresses them to the perimeter of the trough. A Hamilton syringe is then used to deliver ~0.5 µL of previously aliquoted lipid to the center of the trough. At this point the thin film of castor oil also serves as a “piston” to control the surface pressure of the lipid monolayer (Langmuir, 1917). We believe that the castor-oil piston is a desirable but not necessary element of our protocol. Using a trough, rather than smaller, individual wells, facilitates use of the castor-oil piston, and allows one to prepare multiple grids from a single lipid monolayer.

We wash Quantifoil grids by first dipping them into chloroform and then into 95% ethanol. We also apply an additional ~5 nm of evaporated carbon to the top side of Quantifoil grids and then allow these to “age” for at least three days to make the freshly evaporated carbon more hydrophobic. Just before use, we again wash these grids by dipping them into 95% ethanol. We have less experience with C-flat grids, but we have successfully used them as received, i.e.without depositing additional carbon. As is mentioned in the Supplemental Material, in the section “*Issues still to be addressed*”, we prefer to use gold or molybdenum grids to copper grids.

In our current crystallization protocol, we dilute an aliquot of SA to a concentration of 0.2 mg/mL with subphase buffer, and we use 4 ¼L of diluted SA for each grid. After applying SA to the EM grid, the crystals are grown within a humidity chamber. Care is taken to minimize evaporation by placing crushed ice on the tweezers, with the intent to cool the grid slightly below the ambient dew point. Whereas an incubation time of 1–2 minutes appears to be too short to ensure full crystal formation, we observed crystallization to be completed within 10 minutes.

Following an incubation time that, for convenience, is often about half an hour when doing many grids at a time, most of the unbound SA is washed away by placing the grid on top of a 200 µL drop of wash buffer. The composition of the wash buffer is 10% trehalose in 10 mM HEPES (pH 7.5), as before, but with the KCl concentration now reduced to 50 mM. After waiting several seconds, the floating grid is caught with a tweezers and lifted vertically until it separates from the wash drop. As noted further in the Supplemental Material, we believe that this step in the protocol may be the one in which the monolayer crystals of SA are at greatest risk of becoming fragmented or even lost completely. Even when crystals have been severely damaged (at this stage, as we believe), most, if not all, holes in the holey carbon film still remain covered by a monolayer of SA bound to biotinylated lipid.

### Streptavidin crystals are then embedded in trehalose and backed with evaporated carbon

After washing the grids to remove unbound SA, excess trehalose solution is gently “wicked off’ by touching the edge of the grid to a piece of filter paper. The grid then is left on a filter paper with the wet side facing the air, and any remaining liquid on the grid is allowed to dry.

After the trehalose solution has dried, a thin layer (~5 nm or less) of evaporated carbon is deposited on the back side (lipid-tail side) of the EM grid. Grids are placed ~20 cm below a carbon-arc source, and evaporation is done at a vacuum of 10^−5^ torr or lower. We use carbon rods with a tip width of 1 mm (Ted Pella catalog number 62-107), which require less power to evaporate, in order to minimize heating of the trehalose-embedded SA “target”. To further minimize the risk of radiant heating, the carbon rod is heated very rapidly, resulting in breakage of the thin tip and a concomitant flash of evaporated carbon. A cartoon showing the structure of the resulting carbon-backed, trehalose-embedded streptavidin support films is presented in Supplemental Figure S2.

We store the carbon-backed, trehalose embedded SA crystals at room temperature in a sealed container. We prefer to store grids over silica gel that is pink (but not white) to maintain a relatively constant value of humidity.

Just before use, the grid is rehydrated by touching to two successive, 50 µL drops of a solution of 10 mM HEPES (pH 7.5) with 150 mM KCl without trehalose and then left on a 100 µL drop of the same solution for 10 minutes. This same process is repeated a second time, with the intent being to rinse away all remaining trehalose. After that, the grid is further washed with whatever buffer is optimal for the macromolecular sample under investigation.

### Preparation of grids and electron microscopy of 70S ribosomal particles

Ribosomes were purified from E. coli strain MRE600 using sucrose gradient centrifugation, as previously described (Blaha et al., 2000). Ribosome complexes were formed by incubating 1.5 µM deacylated tRNA^Phe^ and 3 µM mRNA of sequence 5‘-GGCAAGGAGGUAAAAUUCUACAAA-3’ (Thermo Scientific) with 0.5 µM ribosomes at 37 °C for 15 minutes in the buffer A: 20 mM HEPES, pH 7.5/ 70 mM KCl/ 6 mM MgOAc/ 1 mM TCEP. The antibiotic spectinomycin (Sigma-Aldrich) at a concentration of 20 µM was added to the pre-formed complex and incubated an additional 10 minutes at 37 °C.

Ribosomes were biotinylated by adding 5-fold excess of biotin-labeling reagent (Solulink Catalog No. B-1007-110) and incubated 20 minutes at room temperature. Excess unreacted biotin was removed by using 1 mL S300 Sephacryl (GE Healthcare) gel-filtration spin column. The column was equilibrated with Buffer A, and usually 50 μL of ribosome complex was loaded on the column and spun on a table centrifuge at 3200 rpm for 1 minute.

Grids were first washed 3 times with 50 µL drops of cold ribosome buffer, and then a 4 µL aliquot of ribosomes was applied at a concentration in the range of 20-40 nM. After incubating for 20 minutes, chilled and in a humidity box to minimize evaporation, as described above for growth of monolayer crystals, unbound ribosomes were washed away by touching to three successive, 50 µL drops of buffer.

To prepare negatively stained specimens, we added 4 µL of 2% uranyl acetate to the lens of buffer adhering to the rinsed grid, which then was mixed by repeated, gentle pipetting while on the grid. This was followed by two cycles in which 3 µL was removed from the grid and 4 µL of 2% uranyl acetate was again added. After the second cycle, the excess uranyl acetate solution that remained was removed by blotting with filter paper. Images were recorded with a Gatan CCD camera on a JEOL 1200 electron microscope.

To prepare cryo-EM samples, the grid is transferred to the Vitrobot tweezers after washing away unbound ribosomes with ribosome buffer. The tweezers and grid were then loaded into the Vitrobot Mark IV chamber, which was previously equilibrated at a temperature of 15 °C and a relative humidity setting of 100%. In order to standardize the volume of liquid on the grid before blotting, excess liquid was first wicked off by touching the bottom edge of the grid with filter paper, which was brought in through a side port of the Vitrobot chamber. Following this, 1.2 µL of sample buffer was then added to the wet face of the grid. The blotting time used was 5 s, with a force setting of 8, a blotting time of 3 seconds, and zero wait/drain time.

For routine evaluation of cryo-grids, images were recorded with a Gatan CCD camera on a Philips CM 200. High-resolution images were obtained with a Gatan K2 camera on an FEI low-base Titan, using a Gatan cryo-holder. In this latter case, images were recorded as dose-fractionated movies consisting of twenty 300 ms frames, each with an exposure of 1.2 electrons/Å^2^ at the specimen. The pixel size in these images was 1.3 Å, referred to the specimen. The movie frames were aligned and summed with the motion-correction software developed by (Li et al., 2013).

### Data processing

The “2dx” software package (Gipson et al., 2007) was used to unbend the SA lattice in a few of the images. This was done to sharpen the Bragg spots in the Fourier transforms, thereby increasing the resolution at which spots could be detected with a good signal-to-noise ratio.

In all other cases, Fourier filtration of Bragg peaks was used to remove the image of the SA crystal, without unbending, before particles were boxed for further analysis. To do this, a script, available upon request, was written in MATLAB to identify all pixels where the magnitude of the Fourier transform of an image was higher than a user-defined value. The Fourier-transform magnitudes in such pixels, plus those in several adjacent pixels, were replaced by the average value of the surrounding background, and the phases were replaced by random values. The threshold value was adjusted manually, while looking at a display of the Fourier transform of an image, until all visible diffraction spots were removed.

Candidate ribosome particles were automatically boxed with a software tool provided in RELION (Scheres, 2012). Images, with candidate particles outlined, were edited manually, using the BOXER tool provided in EMAN (Tang et al., 2007) to remove initial candidates that were obviously aggregates or other undesired material. Three-dimensional classification of particles and subsequent refinement again used tools provided in RELION.

## RESULTS

### Characterization of the carbon-backed SA monolayer crystals

Obtaining monolayer crystals by the on-grid technique is quite reliable and reproducible. On the basis of our current work using LEGINON for automated data collection (Suloway et al., 2005), we estimate the success rate to be well over 90 percent for getting images in which single SA crystals cover the full field of view. The estimated ice thickness was the only criterion used when selecting areas to be added to the data-collection queue, and no attempt was made to decide, in advance, whether the holes contained SA crystals. Nevertheless, until the monolayers have been backed with evaporated carbon, our experience indicates that their crystallinity may still be at considerable risk. More information on factors that we believe can cause preparation of these grids to fail is presented in the Supporting Material.

The fact that these SA crystals remain well ordered after carbon coating, removal of trehalose, and application of a sample has been established by recording high-resolution images. One such image is shown in Figure 1A, and the FFT of this raw, unprocessed image is shown in Figure 1B. In addition to showing clear Thon rings from the evaporated carbon, the FFT also shows many Bragg peaks from the streptavidin monolayer crystal. The Fourier transform of the unbent image is shown in Figure 1C, and an enlarged version is shown in Figure S3. The Bragg spots in the FFT of the unbent image are represented by squares of various sizes, the largest corresponding to an IQ (Henderson et al., 1986) of 1 (estimated phase error of ~4 degrees) and the smallest corresponding to an IQ of 7 (estimated phase error of ~45 degrees). As in this example, Bragg spots with an IQ of 3 (estimated phase error of ~14 degrees) commonly extend to a resolution of 4 Å.

**Figure 1.**
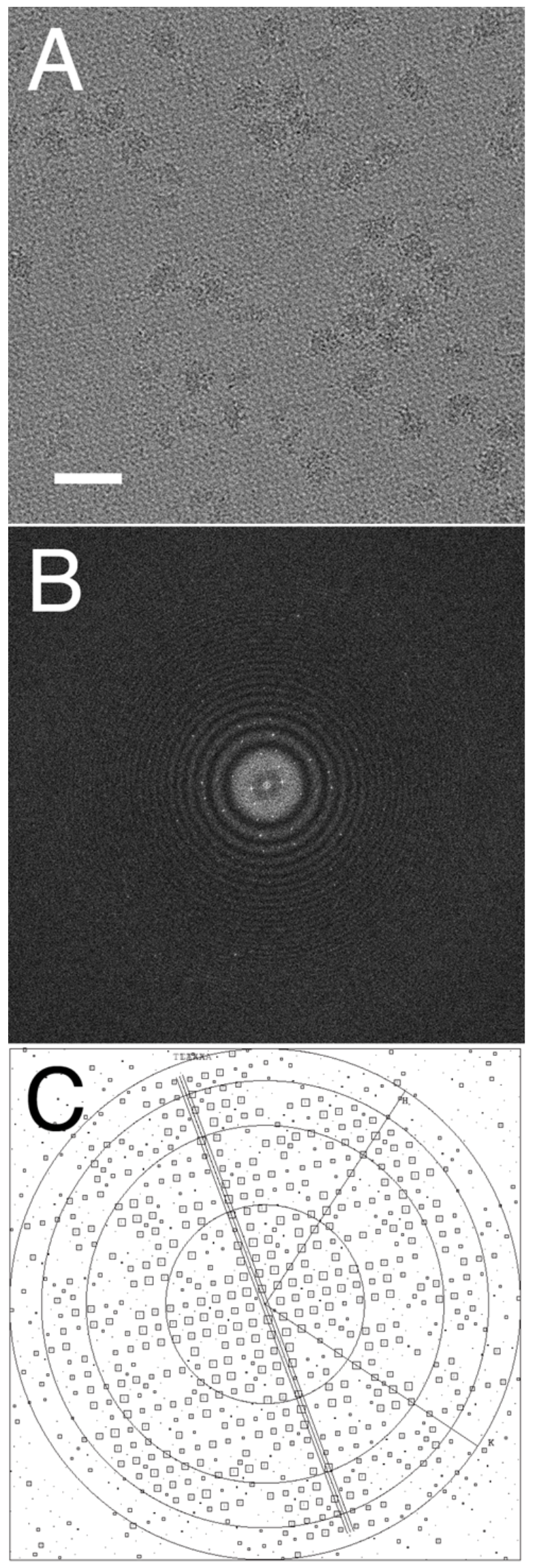
Demonstration that the SA monolayer crystals remain well ordered after trehalose embedding, backing with evaporated carbon, and subsequent removal of trehalose just before use. Panel (A) shows a cryo-EM image recorded with a K2 camera on a FEI Titan electron microscope, operated at 300 keV. In this case, images were recorded as movies consisting of 20 frames, using an electron exposure of 1.2 electrons/Å^2^ per frame. Movie frames were aligned with the software developed by (Li et al., 2013). Panel (B) shows the raw Fourier transform of this image. Panel (C) shows the Fourier transform of the “unbent” image, in which the Bragg spots are displayed as a so-called IQ plot (Henderson et al., 1986), and a larger version of this panel is shown as Figure S3. The 2dx software package (Gipson et al., 2007) was used to unbend the lattice. Indexing of the lattice is indicated by the H and K coordinate system, and the orientation of the specimen-tilt axis, as well as the tilt angle, is also shown in the figure. The signal-to-noise ratio at the reciprocal-lattice points is represented by square boxes of different sizes. Circles are shown at resolutions of 9 Å, 5 Å, 4 Å, and 3.5 Å, respectively. Numerous diffraction spots with IQ values of 4 or less (indicating an expected phase error of 22 degrees or less) are seen out to a resolution of better than 4 Å.

Finally, these grids are easy to prepare and use. The incubation time required to grow streptavidin-monolayer crystals by the on-grid technique is much shorter than the ~2 hours or more that has been recommended for crystallization by the micro-well technique (Wang and Sigworth, 2010). While we often use the SA support films within a week after first making them, we have confirmed that the SA lattice shows no signs of deterioration when stored for a month. We thus expect that the room-temperature shelf life will prove to be much longer than that.

### Use of randomly biotinylated 70S ribosome particles as a test specimen

Figure 2B shows that a high density of ribosome particles is obtained for the conditions of sample concentration and subsequent grid-washing described in the Materials and Methods section. In a corresponding control experiment, using ribosomes that had not been biotinylated, few ribosomes remain on the support film, as is shown in Figure 2A. We thus conclude that the biotin-binding functionality of streptavidin is retained throughout the process of trehalose embedding, carbon evaporation, and storage.

**Figure 2.**
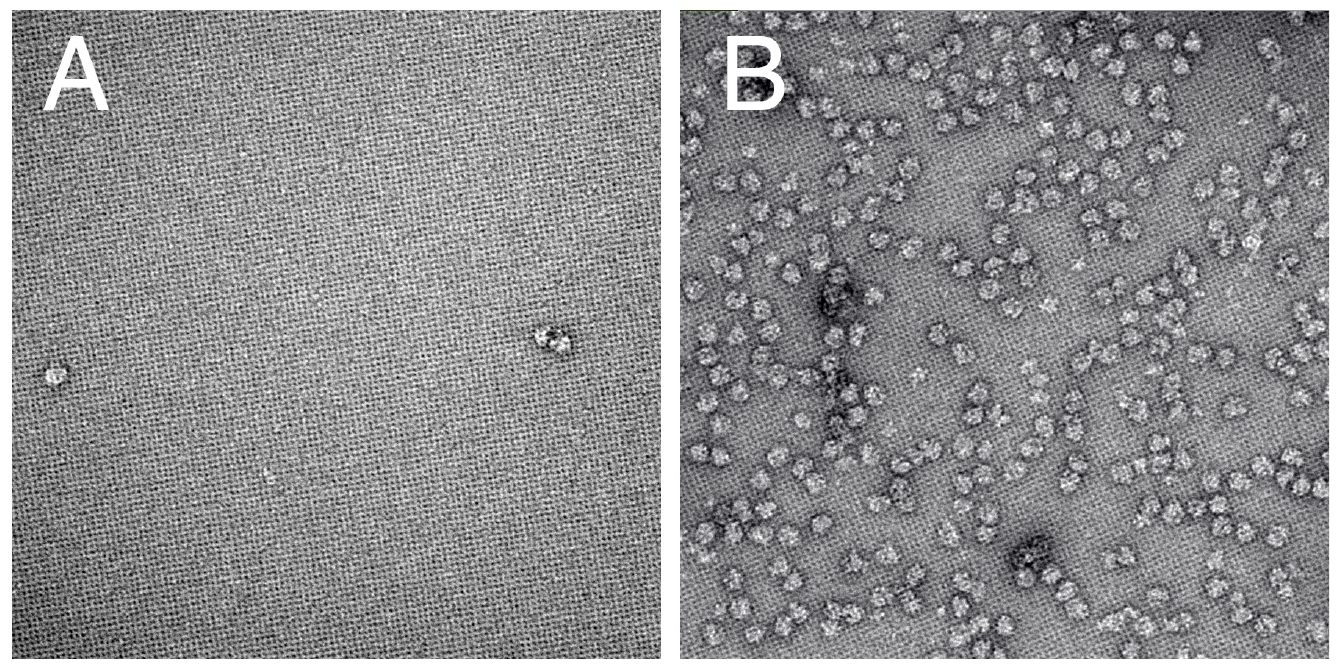
Images demonstrating the effectiveness of binding randomly biotinylated 70S ribosomes to the SA monolayer-crystal support film. (A) This panel shows that only a sparse density of ribosomes was obtained in the control experiment, for which the ribosomes were not biotinylated. (B) By comparison, a uniform, high density of ribosomes was obtained when the ribosomes were biotinylated.

A set of 1497 movies were recorded in order to test the use of SA support films for singleparticle data collection. After removing the SA background from these images, candidate ribosome particles were boxed automatically. This raw set was edited manually to produce a working data set of 101,213 particles. 3-D classification was then used to obtain structurally more homogeneous subsets of the working set.

Refinement of the largest such subset, consisting of 52,433 particles, produced the density map shown in Figure 3A The overall resolution of this map is estimated to be 4.0 Å, based on the point at which the “gold standard” FSC curve, shown in Figure S4, falls to a value of 0.143. Focused refinement, based on the 50 S subunit, improved this only slightly, to just under 3.9 Å. An example of the quality of the map in the interior of the large subunit is shown in Figure 3B.

**Figure 3.**
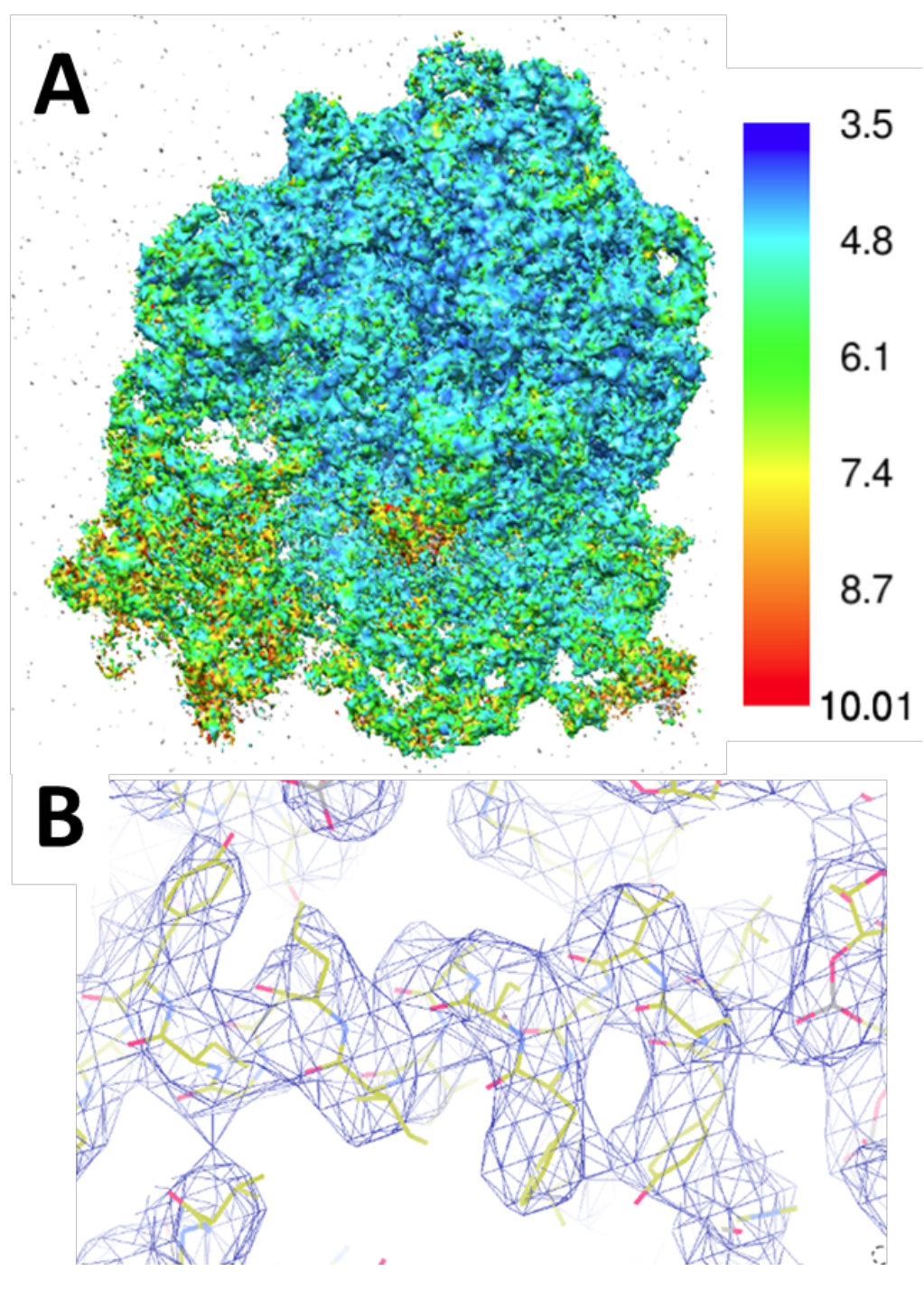
Density map of E. coli 70S ribosome particles obtained when using SA monolayer crystals as a support film. (A) Surface representation of the map, color-coded by the local resolution, with the color scale shown to the right. (B) Example of the map in the best region, with an atomic model of the structure fitted to the map.

A plot of the Euler angles assigned during refinement is shown in Figure 4. These assignments are not distributed uniformly over the surface of the unit sphere, nor do we believe that they ought to be. As is shown in Figure S5 of the Supplemental Material, for example, a significant fraction of the surface of 70 S particles is made up of RNA. Since no biotinylation of RNA can occur, we do not expect particles to bind in the corresponding orientations. In practice, the areas of heaviest assignment of Euler angles correlated roughly, but not always precisely with orientations where there are lysine residues that can be biotinylated. In addition, large portions of the Euler angle plot in Figure 4 have only one or two particles that were assigned to each point. As described by results shown in Figure S6 in the Supplemental Material, we are currently investigating whether this low-frequency, uniform assignment of orientations may reflect errors that are occasionally made in the process of projection matching, rather than non-specific binding.

**Figure 4.**
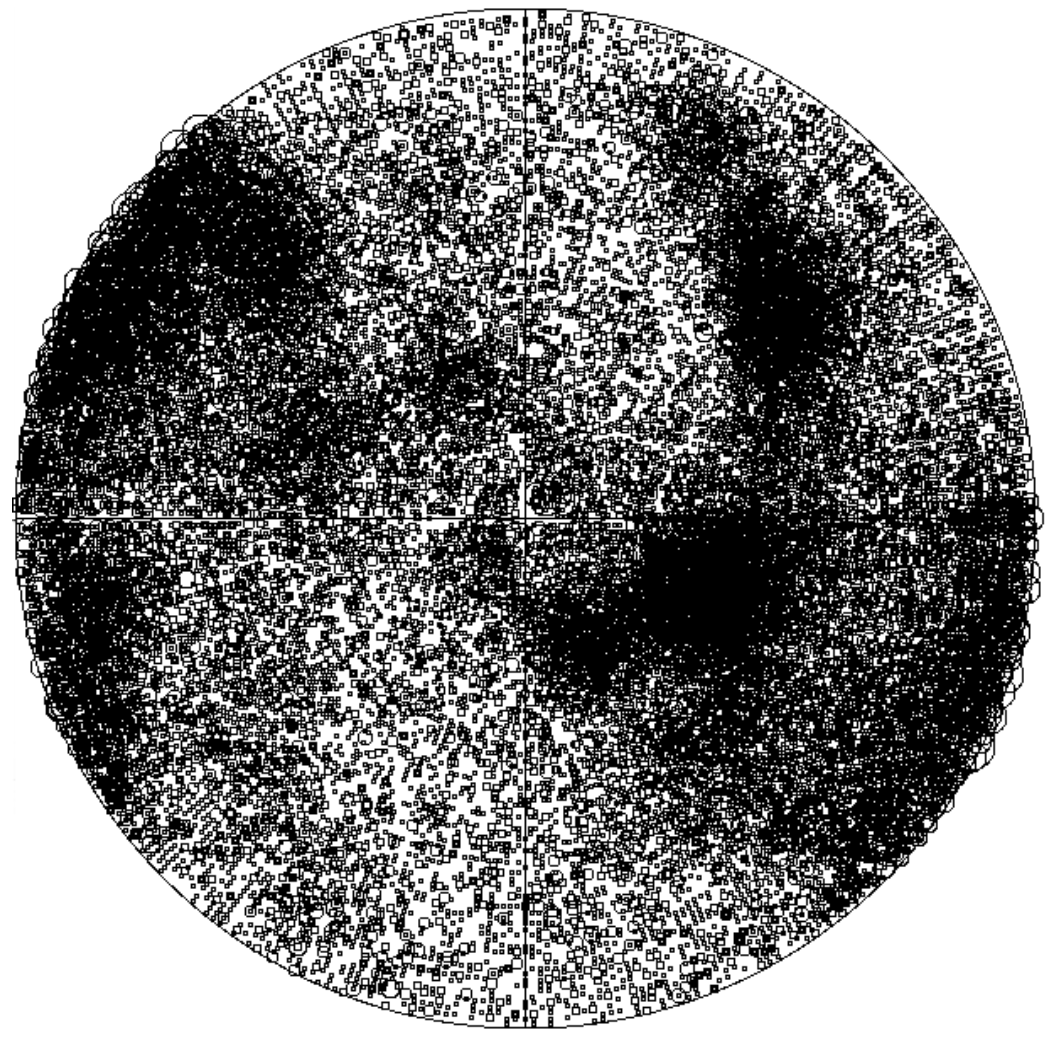
Plot of the angular orientations assigned to 52,433 ribosome particles during refinement of the 3-D density map. Assignments are confined mainly to broad, continuous patches. The remaining particles are distributed quite uniformly, but usually with only one or two particles assigned per point on the unit sphere. As is described in Results, and again in the Discussion, biotinylation (and thus binding) is not expected for those parts of the surface that are made up of RNA. We are currently investigating whether this low-frequency, uniform assignment of orientations might not be accurate, but rather may reflect major errors that are occasionally made in the process of projection matching.

## DISCUSSION

### Streptavidin affinity-grids can be used to obtain high-resolution structures

The resolution of the 3-D density map that we obtained for *E. coli* 70 S ribosomes, using SA monolayer crystals as a support film, is in line with that of maps obtained previously by others, using continuous carbon as a support film. A map at a somewhat lower resolution of ~5.1 Å was obtained by (Bai et al., 2013) for *T. thermophilus* 70 S ribosomes, but from a much smaller data set consisting of 15,202 particles. On the other hand, significantly higher-resolution maps have been obtained, this time for *E. coli* 70 S ribosomes, when using much larger data sets. A 3.0 Å map was obtained from 164,353 particles by (Brown et al., 2016), and an even higher-resolution structure was obtained by (Fischer et al., 2015) from 417,201 particles.

It is natural to be concerned that the periodic motif of the SA crystal might interfere with alignment and assignment of Euler angles for boxed particles. In principle, this should not be a problem because this background is easily removed by Fourier filtration when it is no longer of value to have it there (Han et al., 2012; Wang et al., 2008). Our success in obtaining a high-resolution density map of the 70 S ribosome particle confirms that removal of the periodic SA motif by Fourier filtering was as effective as it was expected to be.

Another concern is that nearly irreversible affinity binding could, in principle, trap the structure of a large complex in one or more off-pathway states. This could happen if some biotin residues are normally inaccessible for binding, for example those that are coupled to lysine residues within deep indentations of the surface. Transient fluctuations of the structure might nevertheless bring such biotin residues sufficiently far out that they would have access to a streptavidin binding site. We can again say that our success in obtaining a high-resolution density map rules out the hypothesis that the trapping of non-native structural variants contributes significantly to the structural heterogeneity of the data set.

### Random biotinylation can result in particles adopting many orientations

One of the major concerns about using affinity grids is that particles might be bound in a preferred orientation with respect to the support film. Chemical biotinylation of one or a few lysine residues per particle, used here, addresses this concern by placing affinity ligands as uniformly as possible over the surface of the particle.

Although the distribution of particle orientations in our case is clearly not uniform, there is sufficient coverage to adequately sample the 3-D Fourier transform of the ribosome particle. Fortunately, not all particle orientations need to be presented for the resulting 3-D structure factors to still be sampled completely. In fact, a complete sampling of Fourier space already occurs for rotation of an object about a single axis. In that case, the location of all projection directions is restricted to a great circle on the unit sphere. The Euler-angle distribution shown in Figure 4 is actually far more complete than that.

As was indicated in Results, the distribution of surface locations where there are potential biotinylation sites agrees only partially, but not completely, with the pattern of densely populated Euler angles. The reasons for there being discrepancies can be complex, and the explanations, once determined, may not be generalizable. Variations in reactivity with the biotinylation reagent (which could be determined by mass spectroscopy of peptide fragments) could, for example, be one factor. Flexibility of protein subunits that are biotinylated is very likely to be another factor. In the latter case, flexible tethering of particles to the SA lattice might allow them to search locally for nonspecific, but preferred binding interactions that would, otherwise, be too weak to produce significant binding.

In addition, large portions of the Euler angle plot in Figure 4 have only one or two particles that were assigned to each point. As described in the Supplemental Material, we are currently investigating whether an almost uniform background distribution, in which there are only one or two particles at a given angle, may reflect errors that are occasionally made in the process of projection matching, rather than binding that was not mediated by biotinylation-i.e. nonspecific binding.

### Significant benefits are expected from using streptavidin affinity-grids

Affinity support films in general provide a number of features that are expected to be helpful for preparing cryo-EM grids. These include the fact that particles are bound at a single plane (single Z-height, for untilted specimens), which is important for CTF correction at high resolution. In addition, tethered particles are prevented from interacting with the air-water interface, provided that the aqueous sample is not too thin.

Affinity binding is, in general, expected to leave structures in a close-to-native state. Streptavidin affinity grids have the special feature that binding of the biotin ligand is essentially as strong as a covalent bond.

In addition, the background image of the SA lattice might serve as a fiducial to aid alignment of movie frames, and even sub-regions of frames. The background image of the SA lattice might
also serve to track changes in tilt orientation that occur during the course of recording dose-fractionated images.

### Value is added by the long shelf-life feature of these affinity grids

By separating the step when these affinity grids are made from the time when they are actually used, the process of making cryo-specimens becomes little more complicated than it is for using just holey-carbon grids. It is true, of course, that the process of making these affinity grids is still somewhat time consuming. It thus is very helpful that it is possible to make enough of them in advance that the supply does not run out in the middle of a session of freezing specimens.

The step of preserving SA monolayer crystals over open holes required some optimization of the trehalose concentration, in combination with the blotting technique (i.e. wicking from the edge of the grid), used to leave a residual volume of solution to dry. The requirement for optimization is not too surprising in light of the fact that millimeter-scale droplets of aqueous trehalose can leave complex patterns of convection (colloidal-particle deposition), thickness, and stress (cracking) after drying by evaporation (Grant and Grigorieff, 2015). While we have found the protocol described here to be very reliable and simple to use, it is likely that other variations may work equally well. Indeed, spin-coating may be especially well-suited as a way to spread a thin film of trehalose (Abazari et al., 2014).

Trehalose is known to have superior properties for room-temperature preservation of freeze-dried phospholipid vesicles and proteins (Zhou, 2008). It has also been used extensively to preserve the high-resolution structure of thin protein crystals adsorbed to a carbon support film, a few examples of which are (Hebert et al., 1997; Leong et al., 2010; Tang et al., 2007). We note that other hydrophilic solutes might also work as well as does trehalose-see, for example, section 6.4 of (Glaeser et al., 2007).

## ACKNOWLEDGMENTS

This work has been supported in part by NIH grants R01 GM083039, P01 GM051487, and R01 GM065050.

